# Cortical compensation for afferent loss in older adults: Associations with GABA and speech recognition in noise

**DOI:** 10.1101/2022.02.24.481839

**Authors:** Kelly C. Harris, Brendan Balken, James W. Dias, Carolyn M. McClaskey, Jeffrey Rumschlag, James Prisciandaro, Judy R. Dubno

**Affiliations:** Department of Otolaryngology-Head and Neck Surgery, Medical University of South Carolina; Department of Psychiatry and Behavioral Sciences, Medical University of South Carolina

## Abstract

Age-related deficits in auditory nerve (AN) function reduce afferent input to the auditory cortex. The extent to which the auditory cortex in older adults compensates for this loss of afferent input, also known as *central gain*, and the mechanisms underlying this compensation are not well understood. We took a neural systems approach to estimate central gain, measuring AN and cortical evoked responses within 50 older and 27 younger adults. Amplitudes were significantly smaller for older than for younger adults for AN responses but not for cortical responses. We used the relationship between AN and cortical response amplitudes in younger adults to predict cortical response amplitudes for older adults from their AN responses. Central gain in older adults was thus defined as the difference between their observed cortical responses and those predicted from the parameter estimates of younger adults. More central gain was associated with decreased cortical levels of GABA measured with ^1^H-MRS and poorer speech recognition in noise (SIN). Effects of central gain and GABA on SIN occur in addition to, and independent from, effects attributed to elevated hearing thresholds. Our results are consistent with animal models of central gain and suggest that reduced AN afferent input in some older adults may result in changes in cortical encoding and inhibitory neurotransmission, which contribute to reduced SIN. An advancement in our understanding of the changes that occur throughout the auditory system in response to the gradual loss of input with increasing age may provide potential therapeutic targets for intervention.

**Significance:** Age-related hearing loss is one of the most common chronic conditions of aging, yet little is known about how the cortex compensates for this loss of sensory input. We measured AN and cortical responses to the same stimulus in younger and older adults. In older adults we found an increase in cortical activity following concomitant declines in afferent input that are consistent with *central gain*. Increased central gain was associated with lower levels of cortical GABA, an inhibitory neurotransmitter, which predicted poorer speech recognition in noise. The results suggest that the cortex in older adults can compensate for attenuated sensory input by reducing inhibition to amplify the cortical response, but this amplification may lead to poorer speech recognition in noise.

## Introduction

Over the past 40 years a substantial body of evidence has demonstrated that, in tandem with peripheral deficits, profound changes occur in neural morphology and excitatory/inhibitory balance in the adult cortex. Results from animal models show that cortical neurons can remap their receptive fields and rescale sensitivity (gain) following acute cochlear and auditory nerve (AN) deficits, and that these changes can occur following hearing loss (Kotak et al., 2005; Sanes and Bao, 2009; Popescu and Polley, 2010; Engineer et al., 2011) and AN deafferentation (Kujawa and Liberman, 2009). Studies of sensory deprivation in hearing impaired and deaf individuals have demonstrated similar changes throughout the auditory neuroaxis, during development and into adulthood (Dietrich et al., 2001; Husain et al., 2011; Boyen et al., 2013; Sharma et al., 2016). However, this adaptive plasticity in the cortex can include seemingly opposing effects across different experiments, animal models, and peripheral manipulations (Seki and Eggermont, 2003; Kotak et al., 2005; Sanes and Bao, 2009; Dong et al., 2010; Popescu and Polley, 2010; Engineer et al., 2011; Kraus et al., 2011; Salvi et al., 2016), and evidence suggests that plasticity associated with typically chronic conditions may not be well modeled by acute manipulations (Seybold et al., 2012). Although age-related hearing loss is one of the most common chronic conditions of aging, and our aging population is growing, little is known about how the brain adapts to this late-onset progressive decline and degradation of sensory input.

Decreased afferent input is hypothesized to contribute to age-related changes in inhibitory neurotransmission, specifically a loss of GABAergic inhibition throughout the central auditory system, which in turn leads to greater cortical response amplitudes in response to weaker afferent input, a phenomenon known as *central gain* (Gutierrez et al., 1994; Burianova et al., 2009; Caspary et al., 2013; Stebbings et al., 2016). Consistent with this central gain hypothesis, several studies have reported that older adults have cortical responses that are as large as or larger than younger adults, despite potential peripheral deficits in older adults. To date, several studies have attempted to characterize this relationship in humans by examining associations between average pure-tone thresholds (pure-tone average, PTA) and estimates of GABA derived from magnetic resonance spectroscopy. However, the results are mixed (Chen et al., 2013; Profant et al., 2013; Gao et al., 2015; Lalwani et al., 2019). To systematically address these equivocal findings, we used a neural systems approach that incorporates neural activity from the AN to provide a more comprehensive estimate of age-related changes in afferent input and their effects on the cortex. We used these measures to model relationships between the periphery and cortex to test the hypothesis that older adults will demonstrate larger responses at the cortex than predicted by AN activity based on responses from younger adults. Moreover, we predicted that this increased central gain would be associated with lower levels of GABA in auditory cortices.

Evidence from animal models of peripheral deafferentation suggests that reduced inhibition and enhancement of cortical responses contributes to deficits in rather than enhancement of complex auditory processing (Resnik and Polley, 2021). In humans, there is emerging evidence suggesting that lower levels of GABA contribute to poorer speech recognition in noise (SIN) and decreased neural distinctiveness between cortical representations of speech versus music (Lalwani et al., 2019; Dobri and Ross, 2021). However, these studies did not account for peripheral differences between younger and older adults in afferent innervation, and it is unclear the extent to which GABA levels were the result of general age-related decreases in inhibitory control or due to a loss in sensory input. Our within-subject design identified an age-related loss in neural activity at the AN and increased responsiveness at the cortex, consistent with central gain, examined the impact of age and age-related hearing loss on sensory-driven plasticity, identified a potential underlying mechanism to these changes (inhibitory neurotransmission), and determined the influence of sensory-driven plasticity on speech recognition in noise.

## Methods

### Participants

Fifty older adults [aged 55 - 86 yr, mean age = 66.4 yr, 41 female] and twenty-seven younger adults [aged 18 - 29, mean age = 23.5 yr, 18 female] were recruited from the Charleston, S.C., community. Inclusion criteria included English as a native language and a Mini-Mental Status Examination score of at least 27. Exclusion criteria included a history of head trauma, seizures, conductive hearing loss or active otologic disease, self-reported central nervous system disorders, use of neuroactive drugs, and contraindications for safe magnetic resonance imaging scanning. Younger participants were required to have pure-tone thresholds ≤ 20 dB HL from 0.25 kHz to 8 kHz. Older adults were included if their hearing loss at or below 4 kHz did not exceed 65 dB HL. As a result, pure-tone thresholds varied in older adults; some had audiometric thresholds that were similar to younger adults, whereas others had mild-to-moderate sloping sensorineural hearing loss. Audiometric thresholds for the test ear (right ear) for both groups are shown in Figure 1. As an estimate of hearing thresholds, the PTA was calculated for each participant, representing the average hearing thresholds in the test ear from 0.5 kHz to 4 kHz. Participants provided written informed consent before participating in this Medical University of South Carolina Institutional Review Board-approved study.

**Figure 1.**
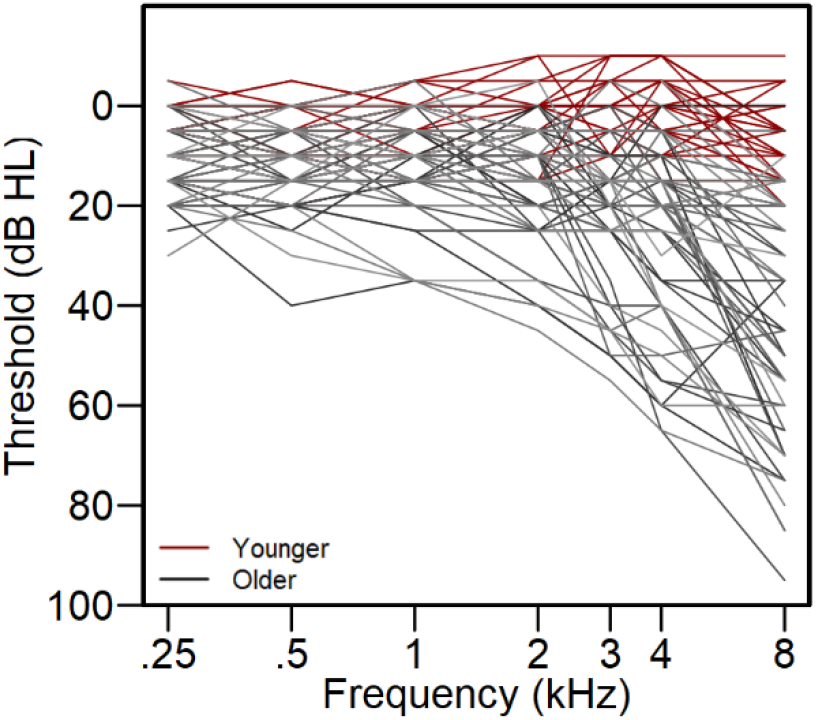
Pure-tone audiograms. Pure-tone air conduction thresholds at audiometric frequencies (0.25 kHz to 8 kHz) for the right (test) ear for younger adults (red lines) and older adults (black lines). Pure-tone thresholds for younger adults were required to be ≤ 20 dB HL at each frequency. Pure-tone thresholds for older adults ranged from 0 to 65 dB HL from 0.25 kHz to 4 kHz.

### Measures of AN and Cortical function

To assess central gain, we examined the amplitude of the AN response and the cortical P1-N1 auditory evoked response elicited in the same participants in response to a 100 μs rectangular pulse. AN responses were estimated from the amplitude of the N1 peak of the compound action potential (CAP), which represents summated activity from the AN. The scalp-measured P1-N1 cortical response amplitude was measured to quantify auditory cortical activity.

#### CAP N1 acquisition and analysis

The N1 of the CAP was elicited by 100 μs rectangular pulses, alternating polarity, presented at 11.1/s through an insert earphone (ER3C; Etymotic Technologies). N1s were recorded in response to stimuli presented at 100 dB pSPL. N1s were recorded in two blocks of 1100 trials (550 of each polarity). N1 responses were recorded using a tympanic membrane electrode (Sanibel) in the test (right) ear, an inverting electrode placed on the contralateral (left) mastoid, and a low forehead grounding electrode. Auditory brainstem responses (ABRs) were simultaneously recorded for reference in identifying Wave I/N1 using a high forehead active electrode, an inverting mastoid electrode (on the right mastoid), and a low forehead grounding electrode. All recordings were collected at a sampling rate of 20 kHz using a custom headstage (Tucker Davis Technologies (TDT), Alachua, FL) connected to the bipolar channels of a Neuroscan SynAmpsRT amplifier (Compumedics USA, Charlotte, NC). Testing was done in an acoustically and electrically shielded room. Participants reclined in a chair and were encouraged to rest quietly for the duration of testing. Participants were allowed to sleep during CAP recording sessions only. Continuous neural activity was analyzed offline in MATLAB (Mathworks Inc., Natick, MA) using EEGlab (Delorme and Makeig, 2004) and ERPLab (Lopez-Calderon and Luck, 2014). Continuous EEG signals were band-pass filtered between 0.150 kHz and 3 kHz. Stimulus triggers were shifted to account for the 1 ms delay introduced by the earphones and the 0.6 ms delay of the TDT digital-to-analog convertor. The filtered data were epoched from −2 to 10 ms and baseline corrected to a −2 ms to 0 ms pre-stimulus baseline (McClaskey et al., 2018). Trials were identified and rejected on the basis of a peak threshold deflection of 45 μV and by visual inspection. Epoched responses for the remaining trials were averaged. N1 peak selection was performed by two independent reviewers and assessed for repeatability across multiple runs. The N1 peak-to-baseline amplitude and peak latency were measured in ERPlab using custom MATLAB functions.

#### Cortical P1-N1 acquisition and analysis

Cortical P1-N1 amplitudes were recorded from a 64-channel Neuroscan QuickCap (international 10-20 system) connected to a SynAmpsRT 64-Channel Amplifier. Bipolar electrodes placed above and below the left eye recorded vertical electro-oculogram activity. Curry 8 software was used to record the EEG signal at a 1 kHz sampling rate. Participants were recorded while passively listening to a 100 μs 100 dB pSPL alternating polarity click with a 2000 ms inter-stimulus-interval (20 ms jitter). The click stimulus was identical to that presented for the CAP but at a slower rate. Two sessions of 200 clicks were recorded from each participant. Continuous EEG data were processed offline using a combination of EEGLab (Delorme and Makeig, 2004) and ERPLab (Lopez-Calderon and Luck, 2014). The recorded EEG data were down-sampled to 0.5 kHz, bandpass filtered from 1-30 Hz, re-referenced to the average of all electrodes, and corrected for ocular artifacts using independent components analysis. Individual trials were then segmented into epochs around the click-onset (−100 ms to +500 ms) and then baseline corrected (−100 ms – 0 ms). Any epochs contaminated by peak-to-peak deflections in excess of 100 μV were rejected using an automatic artifact rejection algorithm. For each participant, epoched data were averaged across trials to compute the average waveform (event-related potential – ERP). ERPs were analyzed in a cluster of frontal-central electrodes, F1, FZ, F2, FC1, FCZ, FC2, C1, CZ, C2 (e.g., Tremblay et al., 2001; Narne and Vanaja, 2008; Harris et al., 2012; Dias et al., 2018). The latencies of the first prominent negative peak (C1/N50), first prominent positive peak (P1), second prominent negative peak (N1), second prominent positive peak (P2), and third prominent negative peak (N2) were recorded. These latencies were used to compute temporal intervals for the automatic detection of the P1, N1, and P2 peak amplitudes and latencies in each of our channels of interest, using custom MATLAB scripts. P1 was defined as the maximum positive peak between C1/N50 (or from stimulus onset if no C1/N50 was detected) and N1. N1 was defined as the maximum negative peak between P1 and P2. P2 was defined as the maximum positive peak between N1 and N2. The P1-N1 amplitude was then calculated as the difference between the positive peak amplitude of P1 and the negative deflection of N1 amplitude, to yield a single cortical peak amplitude for each participant. If a P1 or N1 peak was not found in a channel between their respective latency windows defined from the average waveform across all of our channels of interest, then that channel was omitted from the computed average of each component characteristic (peak amplitude and peak latency). As with previous studies, this approach allowed for automatic peak picking across channels, reducing the risk of human error, while more accurately measuring components by not considering those channels that failed to exhibit an identifiable component, likely as a result of noise from poor impedance (e.g., Anderer et al., 1996; Dias et al., 2018).

### ^1^H-MRS acquisition and processing

Structural magnetic resonance imaging (MRI) scans and ^1^H-MRS data were acquired for a subset of participants with CAP and Cortical responses, N=65 [20 younger (mean age = 23, 15 female), 45 older (mean age = 66, 32 female)]. A Siemens Trio and Prisma 3 T scanner was used to collect images. The same 32-channel head coil was used with both scanners. A structural scan was first taken for voxel placement and tissue segmentation (TR = 5000 ms, TE = 2.98 ms, flip angle = 4°, 176 slices with a 256 × 256 matrix, slice thickness = 1.0 mm, and no slice gap). Following placement of saturation bands 1 cm away from voxel faces and shimming via FASTESTMAP, ^1^H-MRS estimates of GABA were acquired. Due to a scanner upgrade and subsequent change in protocol, a subset of ^1^H-MRS were acquired using a MEGA-PRESS, N=43 [14 younger (mean age = 23, 11 female), 29 older (mean age = 66, 21 female)] sequence with the following parameters (TR = 2000ms; TE= 68 ms, number of averages =150) (Mullins et al., 2014) and a subset using the HERMES sequence, N = 22 [6 younger (mean age = 21, 4 females), 16 older (mean age = 67, 11 females) with the following parameters (TR = 2000 ms; TE= 80 ms, number of averages = 320) (Chan et al., 2016). Unsuppressed water spectra were co-acquired for each sequence and voxel location. ^1^H-MRS data in all sequences were acquired from a 30 mm x 25 mm x 25 mm voxel in our region of interest in left auditory cortex (see Figure 2A). A second voxel was placed in a control region to test the specificity of our effects for auditory cortex, for MEGA-PRESS data a voxel was placed in posterior parietal cortex (Figure 2B), and for HERMES data in occipital cortex (Figure 2C). Each voxel was placed by a trained technician and oriented to avoid including any portion of the skull and ventricles. The auditory cortex voxel was centered on Heschl’s gyrus. The parietal control region was placed above the parietal-occipital fissure, and the visual cortex voxel was centered on primary visual cortex between the parietal-occipital and calcarine fissures. HERMES and MEGA-PRESS offer similar reliabilities for measures of GABA (Prisciandaro et al., 2020).

**Figure 2.**
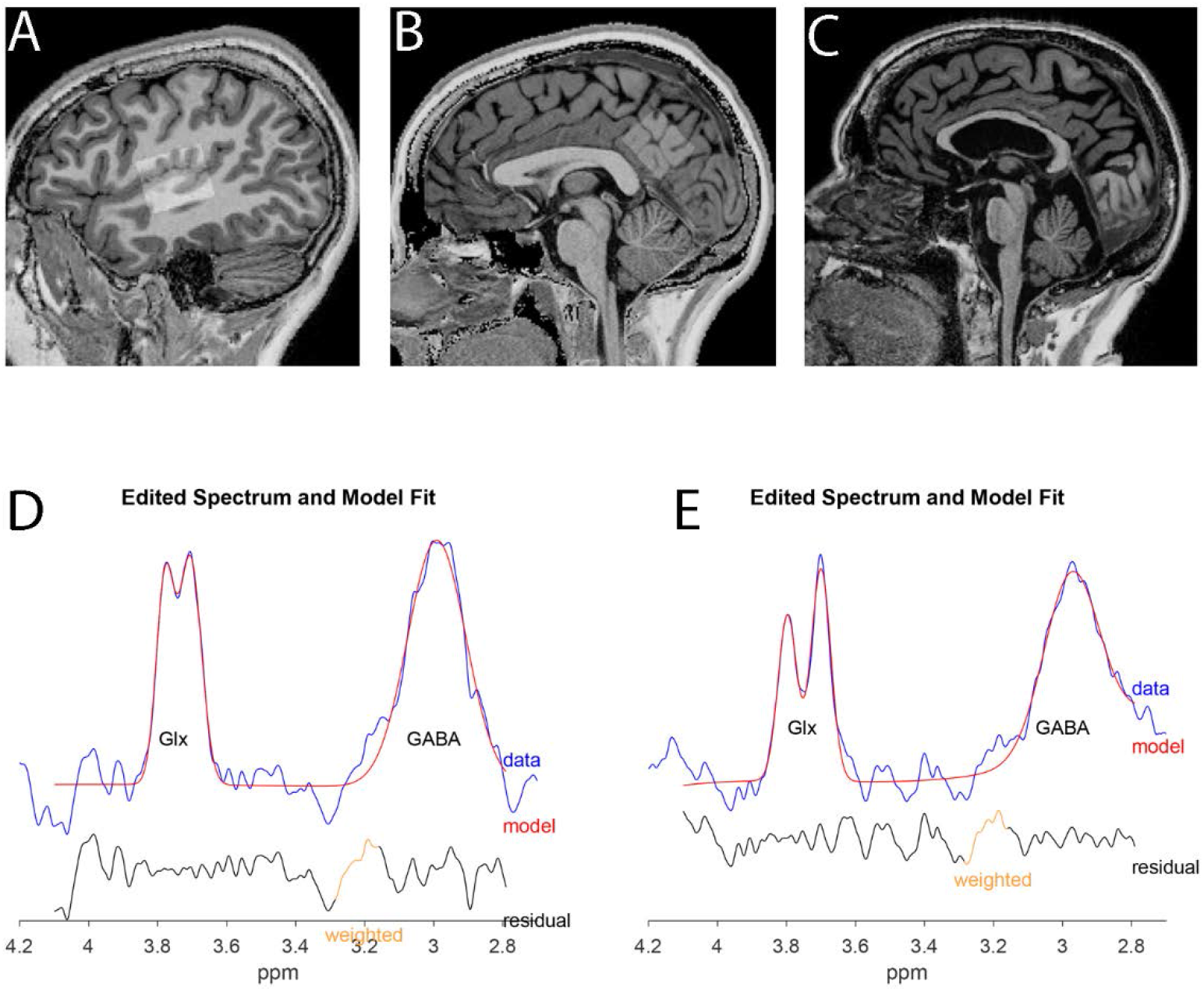
^1^H-MRS voxel placement and GABA spectrum. Sample auditory cortex (A), parietal lobe (B), and occipital lobe (C) voxel placement. Samples were selected from one younger (A) and two older participants (B,C). Note the difference in atrophy between participant A, B, and C and therefore voxel composition across participants. Example of fitted MEGA-PRESS (D) and HERMES (E) from the auditory voxel. The GABA-edited spectrum is shown in blue (plotted across ppm). Overlaid in red is the model of best fit. Below the plot, the residual between these two is shown in black (residual).

GABA values (MEGA-PRESS and HERMES) were processed using the Gannet MATLAB toolbox (Edden et al., 2014). Only metabolites with fitting uncertainties <20% were retained. Water was quantified from a Gaussian-Lorentzian fit to the non-water suppressed data. Quality-control evaluation of ^1^H-MRS spectra resulted in minor data loss, and the number of cases available for analyses was N=59 [20 younger (mean age = 23, 15 female), 39 older (mean age = 65, 11 female)].

Within-voxel tissue fractions of grey matter (GM), white matter (WM) and cerebrospinal fluid (CSF) were calculated based on automated segmentation in Statistical Parametric Mapping 12 (SPM 12, Wellcome Department of Cognitive Neurology) using a volume mask generated in Gannet. Metabolite concentrations (GABA) were normalized to unsuppressed water (metabolite/water). The amount of GABA in the MRS voxel depends in part on its tissue composition. The composition of tissue types is inevitably heterogeneous because of the large size of the MRS voxel, and this effect is even more pronounced in studies examining populations with atrophy due to age or illness. Each tissue type contains a different amount of GABA. GM, where the majority of GABAergic interneurons is found, contains more GABA than WM (Bhattacharyya et al., 2011). CSF contains negligible amounts of GABA. We analyzed two measures of GABA: GABA, the metabolite/water ratio corrected for within-voxel CSF fraction (Prescot and Renshaw, 2013), and non-tissue corrected GABA level, which is still referenced to water but not corrected for tissue concentrations within the voxel. The non-tissue corrected GABA level provides information about the overall number of GABA-containing cells in the cortex. The GABA concentration represents the abundance of intra- and extracellular GABA relative to the cortical volume.

### Speech recognition in noise (SIN)

We measured SIN using the Quick Speech-in-Noise Test (QuickSIN; Etymotic Research; Killion et al. 2004). QuickSIN was collected in a subset of participants with CAP N1 and P1-N1 data, N = 72 [24 younger (16 female), 48 older (35 female)], of these participants all of the younger participants had measures of GABA and 39 of the 48 older adults had measures of GABA. The QuickSIN materials include five lists of six sentences each, with each sentence containing 5 keywords, for a total of 30 keywords in each list. The noise was a four-talker babble. The six sentences in each list progressively decrease in signal-to-noise ratio (SNR) from 25 to 0 dB in 5-dB steps. Sentences were presented binaurally through TDH-39 headphones at a fixed level of 70 dB HL (with noise level varying according to SNR) using a combination of an Onkyo Compact Disk Player and an Interacoustics Audio Traveler (AA222). We computed the average number of keywords (out of 5) correctly identified at each SNR (25, 20, 15, 10, 5, and 0 dB), and summed the averages, for a total possible correct score of 30. QuickSIN performance is reported as SNR loss (25.5 minus total key words correct out of 30), and lower values represent better SIN.

### Data analyses

Statistical analyses were performed in R using multivariate analysis, general linear model and generalized linear mixed model analyses (lme4 (Bates et al., 2015)). Coefficient estimates (B) and standardized coefficients (β) are reported for each comparison. The Benjamini-Hochberg procedure was used to correct for false discovery rate using p-adjust in R.

#### ERP analyses and central gain

We first examined differences in AN and cortical responses for younger and older adults. We used a general linear model multivariate approach to test for the presence of central gain. If central gain is present, older adults would have significantly reduced AN responses as compared to younger adults, but cortical responses that are similar to those of younger adults. Amplitudes of the N1 of the CAP and the P1-N1 cortical response were the outcome variables, and age group was the fixed factor predictor.

Second, we used separate linear regression models in younger and older adults to test associations between AN responses and cortical responses in younger and older groups. We predicted that in younger adults larger AN responses would predict larger cortical responses. Consistent with the central gain hypothesis, we predicted that AN and cortical responses would not be significantly associated in older adults. We used the regression model for the younger adults to determine the extent to which cortical responses in older adults were larger than predicted by their peripheral AN response amplitudes. We calculated the predicted cortical responses based on the parameter estimates and the betas from the regression model for younger adults using the predict.lm function in R. We then subtracted the predicted P1-N1 response amplitudes from the observed P1-N1 response amplitudes and use this as a metric of central gain for older adults [central gain = observed P1-N1 amplitude – predicted P1-N1 amplitude]. Central gain can only be calculated in older adults because predicted P1-N1 amplitudes are based on the regression model from the younger adults.

We then examined the extent to which central gain for older adults was predicted by individual differences in PTA and GABA using linear regression and model testing. Separate regression models were used to test associations between central gain and GABA in auditory cortex and our control region.

#### ^1^H-MRS analysis

We used independent samples t-tests to identify differences in data collected with MEGA-PRESS versus HERMES. No significant differences in demographics (age, sex), fit measures, or metabolites were identified (p > 0.05), and including acquisition type did not improve model fit when examining relationships across metrics. Therefore, data from the two sequences were combined for subsequent analyses. We used independent samples t-tests to identify age-group differences in fit error, and unsuppressed water amplitude. Associations between GABA and age group, PTA, and SIN were examined using linear regression and model testing. Tissue composition (GM/ GM +WM) was entered as a covariate in each model.

#### SIN analyses

We used linear regression and model testing to examine associations between age group, PTA, SIN, central gain, and GABA. First, in older adults, we examined associations between SIN, central gain, and PTA. SNR loss was the dependent variable, with central gain and PTA entered as predictor variables. Next, in both younger and older adults, we examined associations between SIN, age group, GABA, and PTA. To identify effects of age, age group was entered in the model as the predictor variable. We then tested models with GABA entered separately from each voxel location. Next, we entered PTA as a predictor variable in the model. We used model testing to compare the total amount of variance explained in each of our models.

Sex was included as co-variate in each model. However, sex was not a significant predictor of central gain, GABA, or SIN, and did not improve model fit (p> 0.05) and is therefore not reported here.

## Results

### Click-evoked AN and cortical ERP responses in younger and older adults

AN and cortical response waveforms are shown in Figure 3. Consistent with our prior reports and others (Burkard and Sims, 2001; Konrad-Martin et al., 2012; Anderson et al., 2021; Harris et al., 2021a), age group was a significant predictor of N1 AN response amplitude, with older adults exhibiting significantly smaller N1 AN responses than younger adults. In contrast, P1-N1 cortical responses elicited by the same stimulus resulted in response amplitudes that were not significantly different for older and younger adults (Table 1). Individual response amplitudes for the CAP N1 and the cortical P1-N1 response are provided in Figure 4.

**Figure 3.**
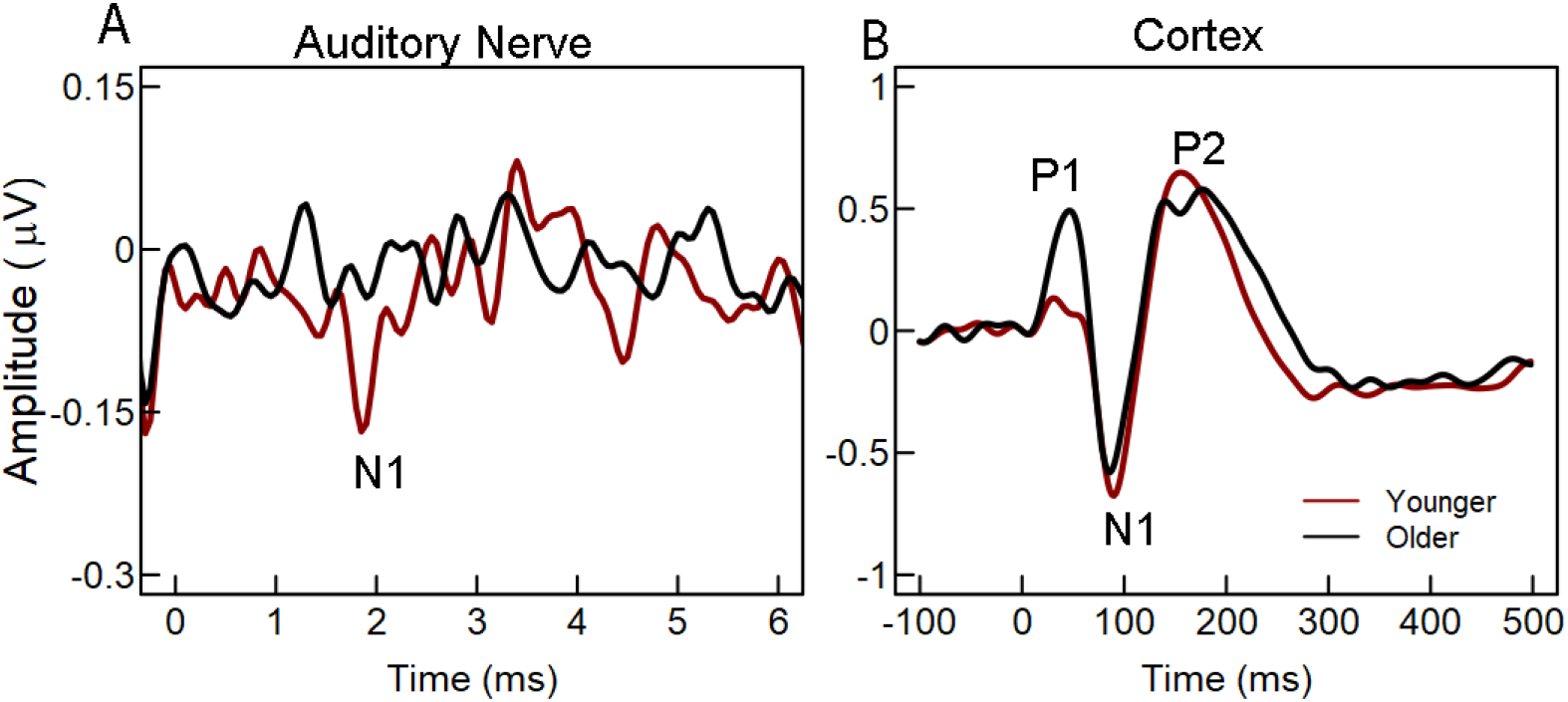
Group average waveforms. **(A)** Group averaged CAP waveforms for younger (red line) and older adults (black line) in response to a 100 dB pSPL click measured from a tympanic membrane electrode. The N1 peak is labeled, indicating the summed population response from the auditory nerve. **(B)** Group averaged cortical P1-N1-P2 waveforms for younger (red line) and older adults (black line) in response to a 100 dB pSPL click measured from a cluster of frontal-central scalp electrodes. The P1, N1, and P2 peaks are labeled. Despite decreases in the CAP N1, cortical responses are similar or larger in older adults compared to younger adults.

**Table 1.** Associations between age group and response amplitudes.

| | $\beta$ | SE | t | p | |
| --- | --- | --- | --- | --- | --- |
| N1 Auditory Nerve Amplitude |  |  |  |  |  |
| Age Group | 0.117 | 0.048 | 2.453 | 0.000 | *** |
| P1-N1 Cortical Response Amplitude |  |  |  |  |  |
| Age Group | 0.275 | 0.166 | 1.658 | 0.101 |  |

Notes: \*\*\*p<0.001.

**Figure 4.**
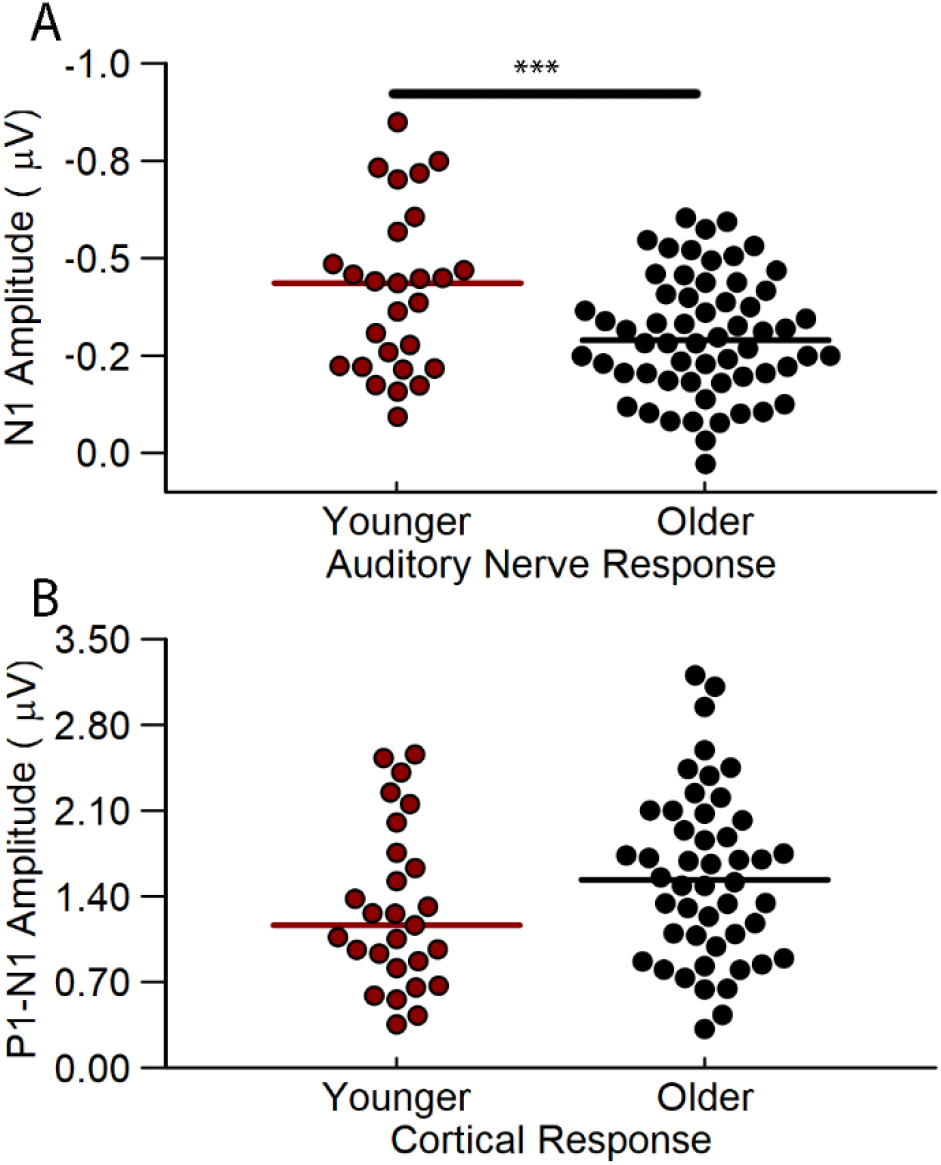
Auditory nerve (N1) and cortical (P1-N1) peak amplitudes. Peak amplitudes for the N1 of the CAP response **(A)** and the cortical P1-N1 response **(B)** for younger (red) and older (black) adults. For comparison, CAP amplitudes are plotted with negative values up, going from 0 to −1 so that in both A and B larger responses are plotted in the positive direction. Median values are marked by a solid line. CAP N1 peak amplitudes were significantly smaller in older than younger adults (p < 0.001 ***). P1-N1 response amplitudes were not significantly different for older and younger adults.

In younger adults, larger CAP N1 responses predicted larger P1-N1 cortical responses [B = −1.42 (SE = 0.56), β = 0.45, t(25) = 2.54, p = 0.017] (Figure 5A), but this was not the case in older adults [B = −0.28 (SE = 0.67), β = −0.061, t(48 = −0.42, p = 0.68] (Figure 5B). We used the predict.lm function to generate predicted P1-N1 amplitudes in older adults, based on the beta estimates from the model in younger adults and the CAP N1 responses of the older adults. Figure 6a plots the predicted P1-N1 response as a function of the observed P1-N1 response for older adults. Observed P1-N1 response amplitudes for most older adults were significantly larger than the predicted P1-N1 amplitude [paired t-test (59) = −4.187, p<0.001] (Figure 6b).

**Figure 5.**
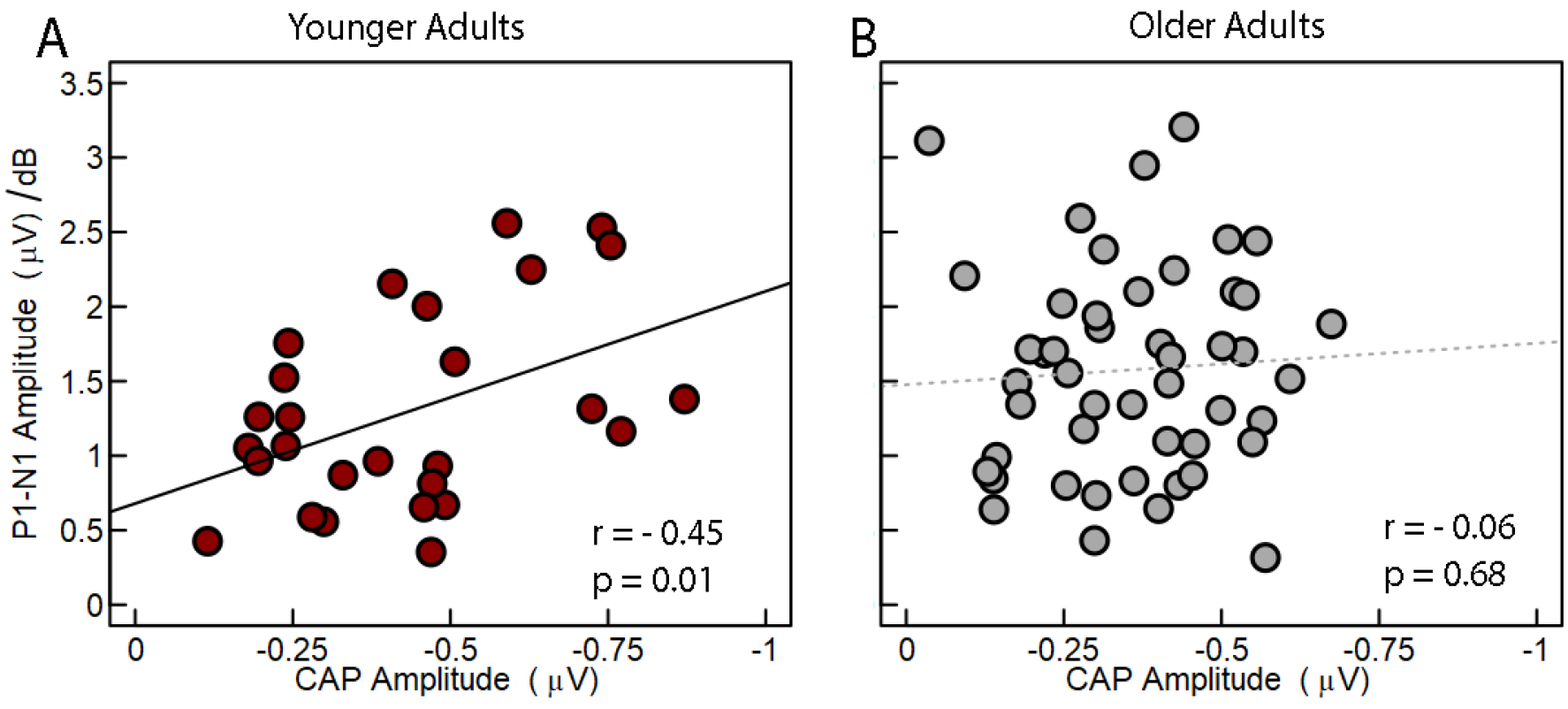
CAP N1 amplitudes predict P1-N1 amplitudes in younger but not older adults. CAP N1 amplitudes plotted relative to P1-N1 amplitude in younger **(A)** and older **(B)** adults. CAP N1 amplitudes are plotted with increasing values (more negative) from left to right. Larger N1 amplitudes were predictive of larger P1-N1 amplitudes in younger but not older adults.

**Figure 6.**
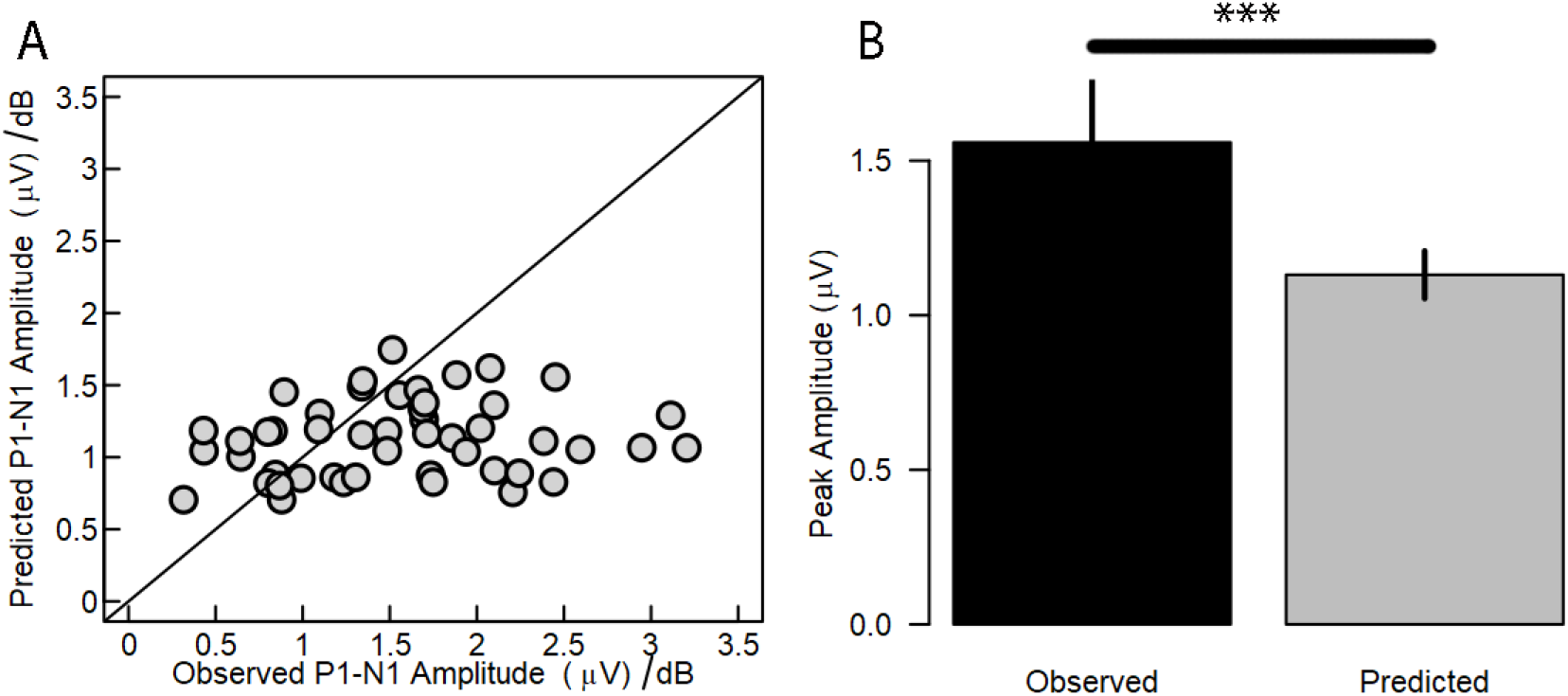
Observed cortical P1-N1 responses in older adults were larger than predicted. Significant associations were observed between the CAP N1 and P1-N1 amplitudes in younger adults (**Figure 4**), and we used that model from younger adults to predict cortical responses in older adults based on their CAP amplitudes. **(A)** Observed P1-N1 response amplitudes plotted against predicted P1-N1 amplitudes for older adults. Data points to the right of the line represent individuals with larger-than-predicted P1-N1 amplitudes. **(B)** Mean observed and predicted amplitudes. Observed P1-N1 response amplitudes for older adults were significantly larger than amplitudes predicted from responses of younger adults (p < 0.001).

### Increased central gain is associated with poorer speech recognition in noise

Consistent with evidence from animal models (Resnik and Polley, 2021), we found that more central gain in older adults was associated with poorer SIN [B = 0.63 (SE = 0.28), β = 0.35, t(38) = 2.28, p = 0.028]. When entered into the model with central gain, PTA was also a significant and independent predictor of SIN (Figure 7, Table 2), and improved model fit [Change R^2^ = 0.12, F(1,37) = 5.66, p = 0.023]. These results suggest that greater central gain in older adults as indicated by their larger cortical responses relative to reduced afferent input may contribute to poorer SIN.

**Figure 7.**
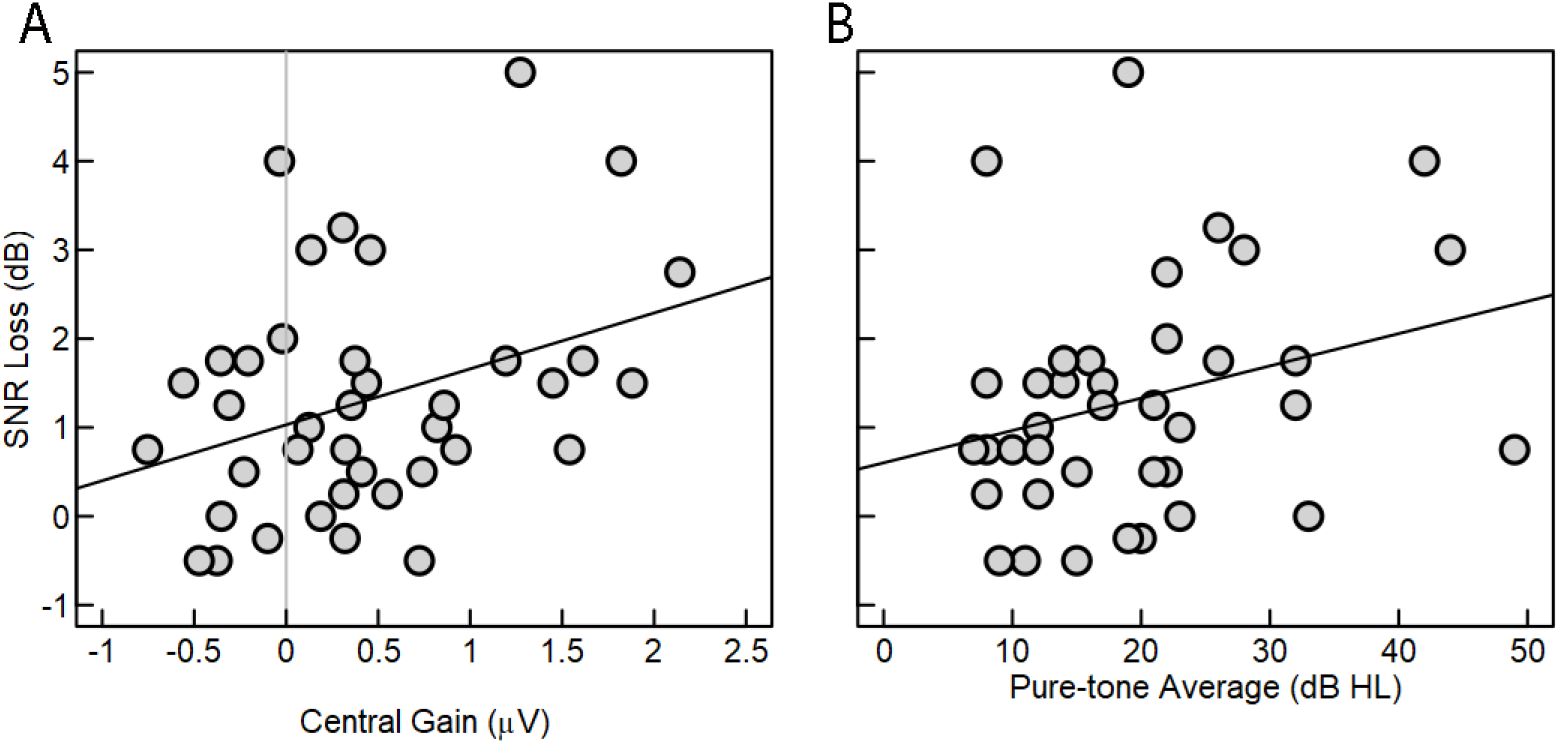
Central gain and PTA predict SNR loss in older adults. Central gain in older adults, plotted as the difference between the observed P1-N1 and the predicted P1-N1 amplitude **(A)**, and PTA **(B)** were significant predictors of SNR loss, with more central gain and more hearing loss contributing to poorer speech-in-noise recognition. Central gain values greater than 0 (indicated by gray vertical line) in panel A represents larger-than-predicted cortical response amplitudes. Solid lines indicate a significant relationship across variables (Table 2). QuickSIN scores are plotted in SNR loss calculated as 25.5 – total correct out of 30 key words, with better speech recognition in noise corresponding to lower values.

**Table 2.** Associations between speech recognition in noise, central gain, and PTA.

| | B | $\beta$ | SE | t | p | |
| --- | --- | --- | --- | --- | --- | --- |
| Model : Predicting QuickSIN scores |  |  |  |  |  |  |
| Central Gain | 0.729 | 0.406 | 0.264 | 2.764 | 0.009 | ** |
| PTA | 0.045 | 0.349 | 0.019 | 2.378 | 0.023 | * |

Notes: \*p<0.05, \*\*p<0.01

### Factors that predict central gain in older adults

We first examined the extent to which PTA predicted central gain. The N1 of the CAP was used as the afferent signal in calculating the predicted P1-N1 for older adults, along with the relationship between the afferent signal and cortical responses in younger adults. Consistent with our previous study (Harris et al., 2021b), individual differences in N1 amplitude were not associated with PTA in older adults (r(38) = −0.05, p = 0.75). PTA was also not a significant predictor of central gain [B = 0.01 (SE = 0.01), β = −0.14, t(38) = −0.99, p = 0.33]. In our second model, GABA (CSF corrected) was entered as a dependent variable. To control for differences in atrophy within the voxel, the amount of gray matter within the voxel was included as a dependent variable. GABA was a significant predictor of central gain, with lower levels of GABA associated with more central gain (Figure 8, Table 3). This association was specific to GABA in Heschel’s gyrus, and not associated with GABA in our control region [B = 0.16 (SE = 0.35), β = −0.09, t(38) = −0.47, p = 0.64].

**Figure 8.**
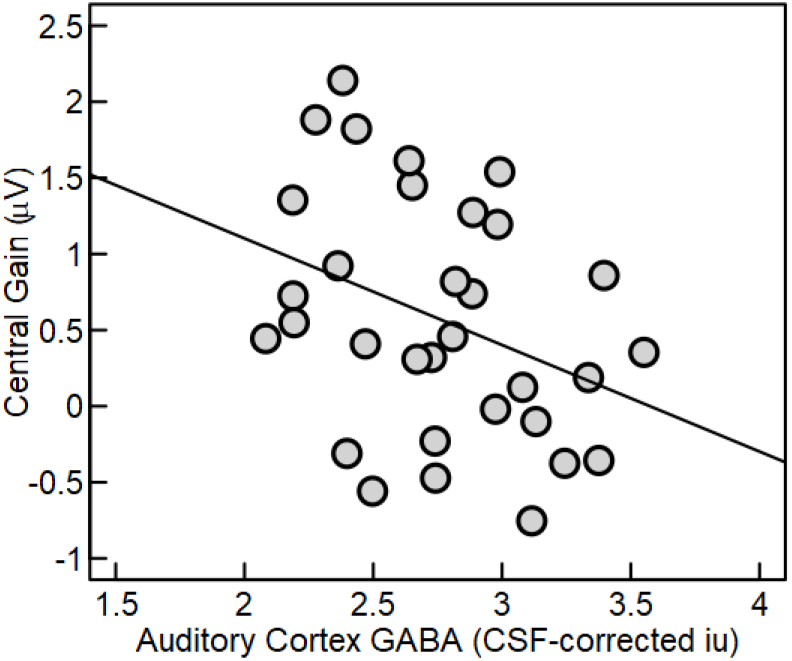
Lower levels of GABA+ in auditory cortex predict greater central gain. Central gain, defined as the observed P1-N1 amplitude minus the predicted P1-N1 amplitude in older adults, plotted as a function of GABA levels in auditory cortex. Solid line represents the significant association between GABA and central gain.

**Table 3.**
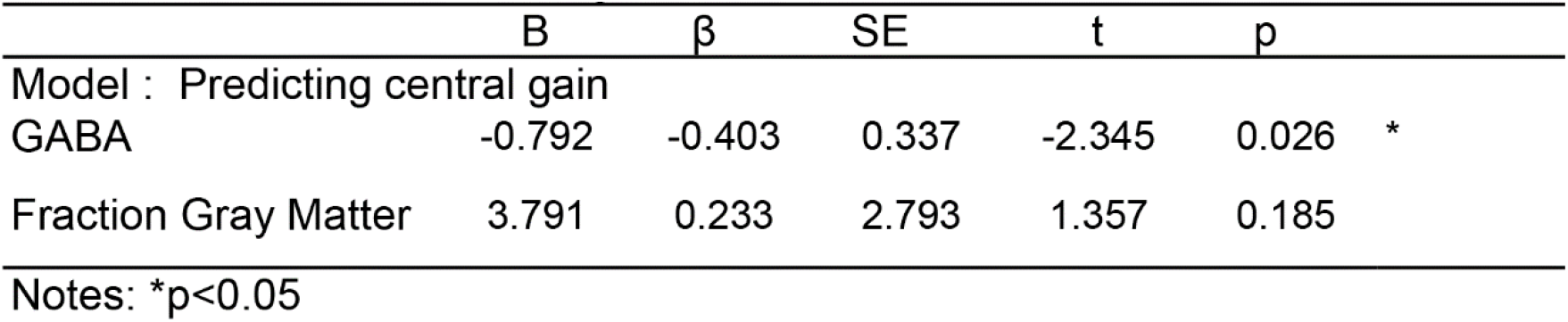
Associations between central gain and GABA.

### GABA: effects of age group, PTA, and cortical region in younger and older adults

Prior to examining effects of age group on GABA we examined several factors that may affect GABA fit, including fit error and unsuppressed water amplitude, and water linewidth. These values did not differ between younger and older adults (p>0.05).

Similar to prior studies, we examined the effects of age on GABA as both non-tissue corrected GABA and CSF-corrected GABA (GABA). These analyses are important as older adults showed greater atrophy within both Heschl’s gyrus [t(58) = 6.32, p <0.001] and our control regions [t(58) =3.55, p < 0.001], calculated as an increase in the CSF fraction of the voxel (decrease in GM and WM). We found a significant effect of age on non-tissue corrected GABA levels in our auditory cortex region, however after including the total fraction of GM and WM within the voxel, total GABA was no longer a significant predictor (Table 4), suggesting that age-related atrophy resulted in lower levels of total GABA in older adults. Estimates of GABA corrected for CSF account for these differences in atrophy, and therefore there was not a significant effect of age on GABA (corrected for CSF) [B = 2.22 (SE = 3.62), β = 0.11, t(58) = 0.61, p = 0.54]. These results highlight two factors that co-occur in auditory cortex with age, atrophy and a loss of GABA.

**Table 4.** Age-group difference in GABA.

| | B | $\beta$ | SE | t | p |
| --- | --- | --- | --- | --- | --- |
| Model 1: Age Group |  |  |  |  |  |
| Age Group | -0.245 | -0.255 | 0.122 | -2.011 | 0.049 * |
| Model 2: Age Group and Gray Matter (GM) |  |  |  |  |  |
| Age Group | 0.053 | 0.055 | 0.127 | 0.416 | 0.679 |
| Fraction GM | 5.145 | 0.569 | 1.201 | 4.28 | 0.000 *** |

Notes: \*p<0.05, \*\*p<0.01,

In our control region, despite significant differences in tissue composition, total GABA [B = 0.05 (SE = 0.11), β = 0.06, t(58) = 0.46, p = 0.65] and GABA [B = 0.08 (SE = 0.12), β = 0.08, t(58) = 0.65, p = 0.52] did not differ across age group. GABA levels were significantly higher in our control region than in auditory cortex [t(58) = −2.12, p = 0.036], even after accounting for differences in SNR and water linewidth across regions. Taken together, older adults exhibited significant atrophy within both auditory cortex and our control region, but only significant differences in total GABA levels in our auditory voxel.

We examined associations between GABA and PTA within older adults and across our sample of younger and older adults. PTA was not a significant predictor of GABA in our sample of older adults in auditory cortex [B = 5.77 (SE = 3.10), β = 0.33, t(38) = 1.86, p = 0.07]. PTA was not a significant predictor of GABA in our control region [B = 2.22 (SE = 3.62), β = 0.11, t(38) = 0.61, p = 0.54]. We next examined whether GABA was predicted by PTA across our sample of younger and older adults. We included GABA as the dependent variable, and PTA and age group as predictors. Again, PTA was not a significant predictor of GABA [B = −005 (SE = 0.01), β = −0.09, t(58) = −0.47, p = 0.64].

### SIN: associations with age group, GABA and PTA

We then used linear regression and model testing to determine the extent to which age group, GABA, tissue composition (GM/GM+WM), and PTA predicted SIN. Models are provided in Table 5. Age group (Model 1) was not a significant predictor of SIN. GABA was a significant predictor of SIN, with lower levels of auditory cortex GABA predicting poorer SIN (Model 2). We then added PTA, which was found to be a significant predictor of SIN in older adults. Both PTA and GABA remained significant predictors of SIN (Figure 9, Model 3), and improved model fit [X^2^(1) =4.57, p = 0.038]. GABA from our control region was not a significant predictor of SIN [B = 0.23 (SE = 0.53), β = 0.06, t(58) = 0.43, p = 0.67], and adding GABA from the control region to the above models did not improve model fit [X^2^(1) =0.56, p = 0.46]. These results suggest that auditory cortex GABA and pure-tone thresholds are each unique predictors of SIN in both younger and older adults.

**Table 5.**
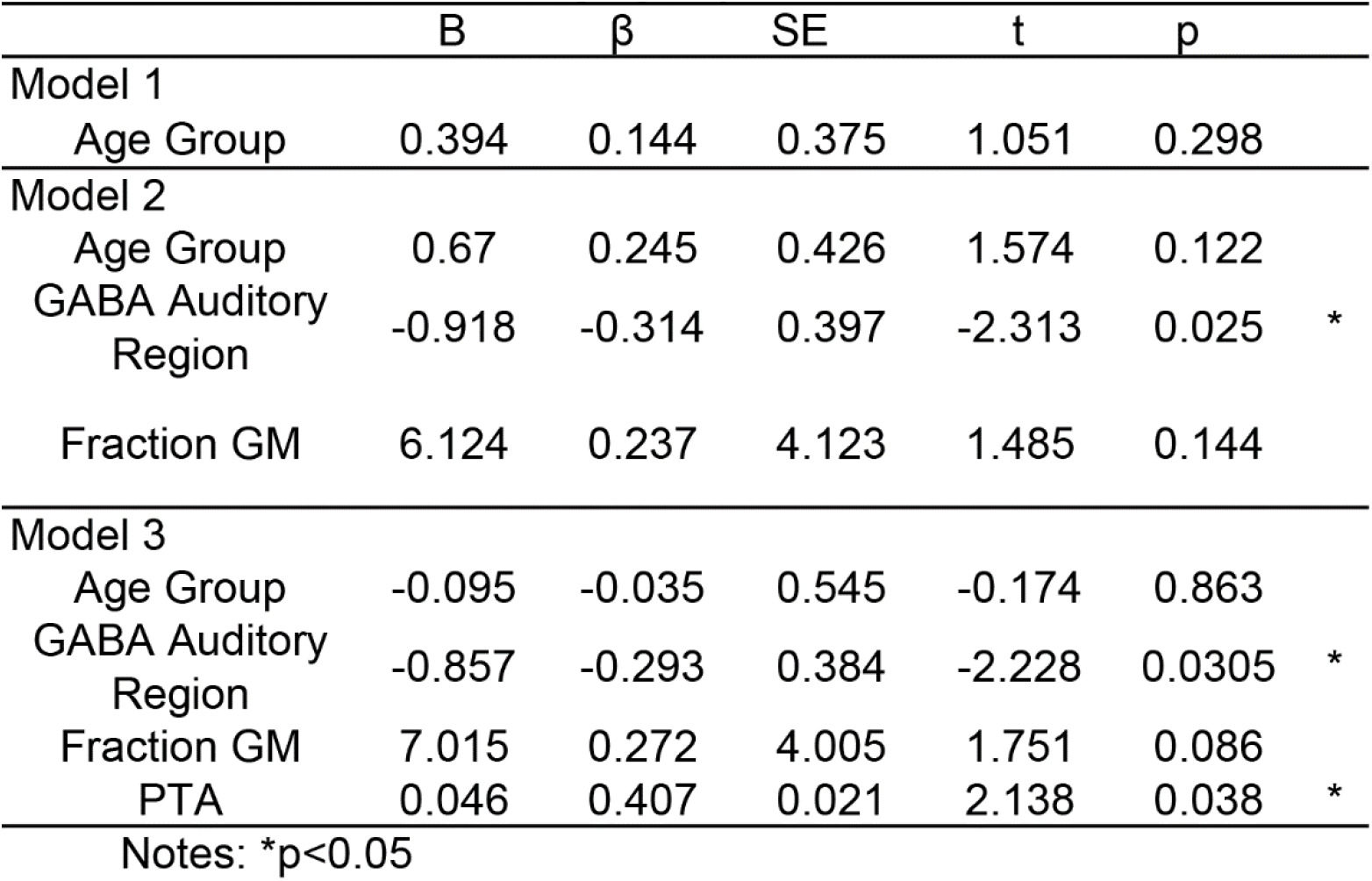
Associations between SIN, age group, GABA+, and PTA.

| | B | $\beta$ | SE | t | p |
| --- | --- | --- | --- | --- | --- |
| Model 1 |  |  |  |  |  |
| Age Group | 0.394 | 0.144 | 0.375 | 1.051 | 0.298 |
| Model 2 |  |  |  |  |  |
| Age Group | 0.67 | 0.245 | 0.426 | 1.574 | 0.122 |
| GABA Auditory Region | -0.918 | -0.314 | 0.397 | -2.313 | 0.025 * |
| Fraction GM | 6.124 | 0.237 | 4.123 | 1.485 | 0.144 |
| Model 3 |  |  |  |  |  |
| Age Group | -0.095 | -0.035 | 0.545 | -0.174 | 0.863 |
| GABA Auditory Region | -0.857 | -0.293 | 0.384 | -2.228 | 0.0305 * |
| Fraction GM | 7.015 | 0.272 | 4.005 | 1.751 | 0.086 |
| PTA | 0.046 | 0.407 | 0.021 | 2.138 | 0.038 * |

Notes: \*p<0.05

**Figure 9.**
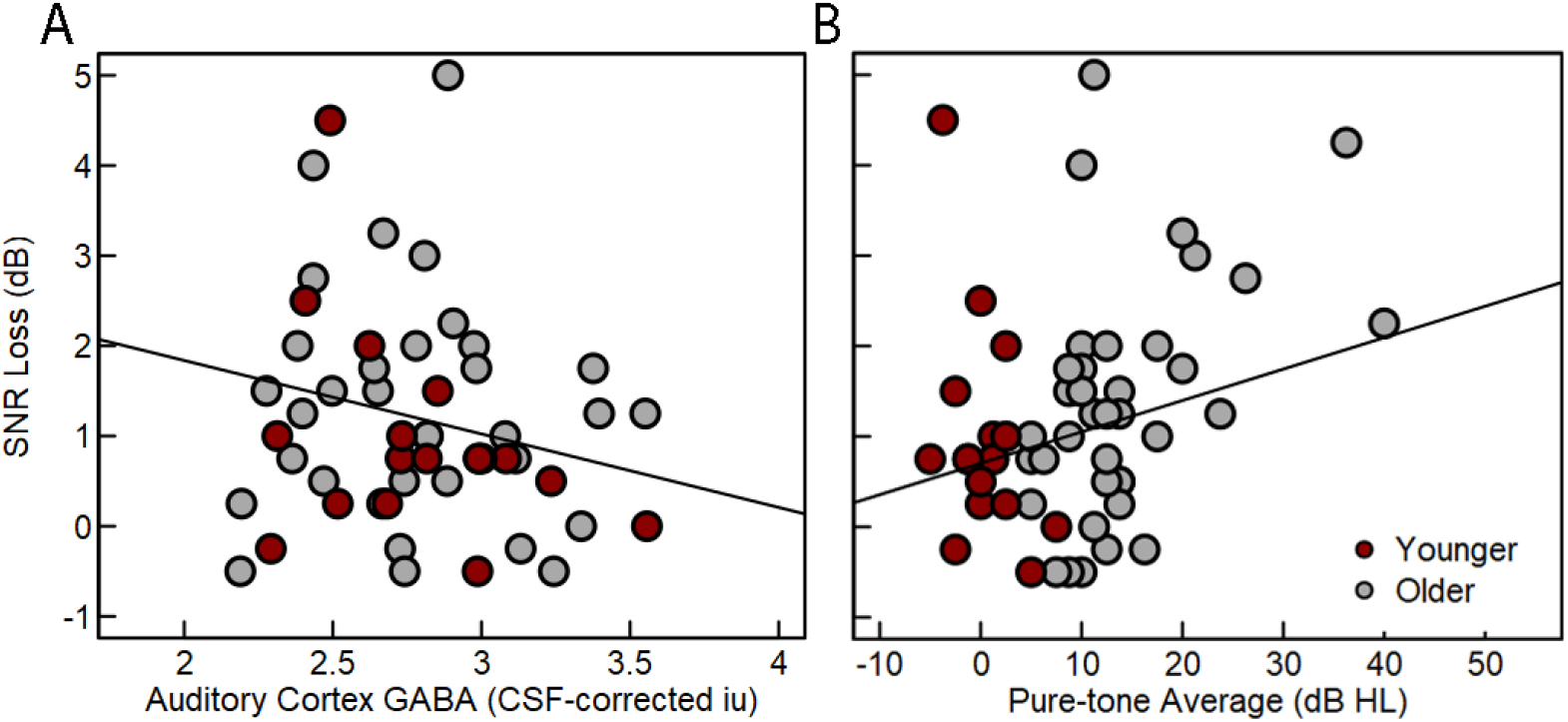
Auditory GABA+ and PTA predicted SNR loss. Lower levels of cortical GABA+ in auditory cortex predicted poorer speech recognition in noise (higher SNR loss) in older (gray circles) and younger (red circles) adults **(A)**. Adding PTA to the model significantly improved model fit, and higher PTA thresholds were associated with poorer speech recognition in noise **(B)**. Statistics are provided in Table 5, solid lines represent significant associations across variables.

## Discussion

The underlying mechanisms and perceptual consequences of sensory-driven plasticity in older adults are largely unknown. Evidence from animal models suggests that both aging and hearing loss result in a slow peripheral deafferentation that propagates throughout the auditory system and contributes to changes in excitation and inhibition at the level of auditory cortex (Caspary et al., 2008). Translating this to older humans, the current study 1) demonstrates central gain in older adults by comparing neural responses from both the AN and cortex, 2) identifies a potential underlying mechanism contributing to this gain, specifically lower levels of GABA, and 3) demonstrates that increasing central gain and lower GABA levels are associated with poorer SIN.

### Central gain: decreased afferent innervation and inhibition contribute to larger cortical responses

Our results are consistent with animal models of central gain and suggest that individual differences in AN afferent input may contribute to changes in cortical encoding. Consistent with our prior work and others (Burkard and Sims, 2001; McClaskey et al., 2018; Anderson et al., 2021; Harris et al., 2021b), we demonstrated a robust decrease in AN response amplitudes for suprathreshold stimulus levels that occurs independently of differences in PTA, suggesting decreased afferent innervation in older adults. Prior studies have examined central gain in the periphery by comparing wave I of the auditory brainstem response to wave V, generated in the midbrain and reported decreased wave I amplitudes relative to wave V amplitudes in older adults (Grose et al., 2019). Extending these results to the cortex, we observed larger-than-predicted cortical responses relative to AN responses in many older adults. These effects were not, however, universal in older adults (Figure 6A), and seem to be driven by changes in cortical levels of inhibition, with decreased afferent inhibition (smaller AN response amplitudes) and lower levels of GABA contributing to larger-than-predicted cortical responses.

### Lack of association with pure-tone thresholds

Changes in AN activity, central gain, and GABA were not associated with individual differences in PTA. It is now well-established that individual differences in PTA are not associated with suprathreshold AN responses in older adults (Burkard and Sims, 2001; Konrad-Martin et al., 2012; Grose et al., 2019; Harris et al., 2021b), and that deficits in AN activity occur in older adults with a range of pure-tone thresholds. To date, associations between PTA and GABA+ in auditory cortex have been equivocal, with one study showing that increased hearing thresholds were associated with lower levels of GABA in older adults (Gao et al., 2015), whereas other studies show no associations (Profant et al., 2013; Dobri and Ross, 2021; Lalwani et al., 2021). The current study examined this association in a relatively large sample of older adults with a wide range of PTAs and found no association between GABA and PTA in older adults or when examining associations across age groups. Central gain was also not associated with PTA. Taken together, these results suggest that standard clinical assessments of hearing thresholds may not be indicative of age-related changes in afferent innervation and sensory-driven plasticity.

### Effects of age group on GABA

Consistent with results from the current study, prior studies often demonstrate an age-related reduction in GABA when examining GABA without correcting for atrophy and tissue composition within the voxel, suggesting that age-related reductions in GABA are due in part to age-related atrophy (Lalwani et al., 2019; Dobri and Ross, 2021). However, age-related atrophy was evident in both our auditory and control regions, yet only GABA from auditory cortex showed an age-related decrease. Effects of age on GABA in sensory cortices appears to be an area where future study is needed, as levels of GABA in visual cortex have been reported to decrease (Simmonite et al., 2019; Chamberlain et al., 2021), and increase (Pitchaimuthu et al., 2017) with increasing age depending on the study, similarly to findings in auditory cortex.

### Relationship to SIN

Greater central gain and lower levels of GABA contributed to poorer SIN. By examining central gain and GABA within the same participants, our results bridge work from animal models that suggest that central gain in the auditory system may disrupt signal in noise encoding (Resnik and Polley, 2021). Here in humans, lower levels of cortical GABA in older adults are associated with poorer SIN and decreased neural distinctiveness (Lalwani et al., 2019; Dobri and Ross, 2021). Moreover, these associations occur independently of the effects of PTA on SIN in both younger and older adults, suggesting that individual differences in GABA not related to age or hearing loss contribute to SIN. This sensory-driven plasticity is thought to reflect a homeostatic mechanism where cortical ensembles strive to maintain levels of neural activity despite a loss of afferent input. Although largely thought to be compensatory in nature, these mechanisms may not work efficiently, especially under complex listening conditions, which could result in poorer auditory processing and poorer speech recognition. Several factors may contribute to associations between central gain and GABA+ and SIN. Animal models suggest that reduced inhibitory control and central gain may disrupt noise encoding and/or contribute to deficits in temporal processing (Gleich and Strutz, 2011; Resnik and Polley, 2021). Future studies are currently needed to identify the extent to which central gain in older adults is associated with deficits in signal encoding and temporal processing.

### Potential target for intervention

Understanding associations between sensory loss, cortical inhibition, neural markers, and behavioral performance may lead to targeted intervention strategies and biomarkers for intervention. For example, in the cognitive domain, greater brain signal variability, measured with EEG, is associated with better cognitive performance and associated with higher levels of cortical GABA. In older adults with poor neuronal variability, pharmaceutically boosting levels of GABA increased neural variability (Lalwani et al., 2021). In the auditory system, central gain occurs without restoring afferent input, and it is unknown how increasing GABA at the cortex and decreasing central gain can account for these remaining deficits and contribute to improved SIN. However, our results suggest that poorer SIN was associated with larger central gain, which reflects amplified cortical activity beyond that predicted by reduced AN function and may therefore be amenable to intervention.

## Summary and Conclusions

In summary, these results demonstrate: 1) Age-related differences in the relationships between peripheral and central auditory responses that are consistent with central gain in older adults, 2) Lower levels of GABA in older adults contribute to central gain, 3) GABA levels do not appear to be driven by PTA or age group, 4) lower GABA levels in auditory cortex in older adults may be dependent on increased cortical atrophy, and 5) Increased central gain, lower levels of GABA, and increased PTA are associated with poorer SIN. Taken together, results suggest that sensory-driven plasticity in older adults may be maladaptive in relation to SIN and highlights a potential factor that contributes to the variation in SIN not predicted by PTA. Several questions remain, including the auditory factors that underlie associations between enhanced cortical responses, lower levels of GABA, and poorer SIN in older adults, and the extent to which altering GABA may ameliorate these deficits and contribute to improved SIN.

## Acknowledgements

We thank the participants of our study. We also thank Lilyanna Kerouac for her assistance with data collection. This work was supported (in part) by grants from the National Institute on Deafness and Other Communication Disorders (NIDCD) of the National Institutes of Health (NIH), R01 DC 014467, R01 DC 017619, P50 DC 000422, and T32 DC 014435. The project also received support from the South Carolina Clinical and Translational Research (SCTR) Institute with an academic home at the Medical University of South Carolina, NIH/NCRR Grant number UL1 RR 029882. This investigation was conducted in a facility constructed with support from Research Facilities Improvement Program Grant Number C06 RR 014516 from the National Center for Research Resources, NIH.

